# Genetic Effects of Welding Fumes on the Development of Respiratory System Diseases

**DOI:** 10.1101/480855

**Authors:** Humayan Kabir Rana, Mst. Rashida Akhtar, M. Babul Islam, Mohammad Boshir Ahmed, Pietro Lio’, Julian M.W. Quinn, Fazlul Huq, Mohammad Ali Moni

**Affiliations:** Department of Computer Science and Engineering, Green University of Bangladesh; Department of Computer Science and Engineering, Varendra University, Rajshahi, Bangladesh; Department of Applied Physics and Electronic Engineering, University of Rajshahi, Bangladesh; School of Civil and Environmental Engineering, University of Technology Sydney, NSW 2007, Australia; Computer Laboratory, The University of Cambridge, 15 JJ Thomson Avenue, Cambridge, UK; Bone Biology Division, Garvan Institute of Medical Research, Darlinghurst, NSW, Australia; Discipline of Pathology, School of Medical Sciences, Faculty of Medicine and Health, The University of Sydney, Australia

**Author notes:** These authors hold joint first authorship of this work (Humayan Kabir Rana, Mst. Rashida Akhtar). Corresponding author: *Email address* (Mohammad Ali Moni).

**Keywords:** Welding fumes, Respiratory system diseases, Chronic Bronchitis, Asthma, Pulmonary Edema, Lung Cancer

## Abstract

**Background:** The welding process releases potentially hazardous gases and fumes, mainly composed of metallic oxides, fluorides and silicates. Long term welding fume (WF) inhalation is a recognized health issue that carries a risk of developing chronic health problems, particularly respiratory system diseases (RSDs). Aside from general airway irritation, WF exposure may drive direct cellular responses in the respiratory system which increase risk of RSD, but these are not well understood.

**Methods:** We developed a quantitative framework to identify gene expression effects of WF exposure that may affect RSD development. We analyzed gene expression microarray data from WF-exposed tissues and RSD-affected tissues, including chronic bronchitis (CB), asthma (AS), pulmonary edema (PE), lung cancer (LC) datasets. We built disease-gene (diseasome) association networks and identified dysregulated signaling and ontological pathways, and protein-protein interaction sub-network using neighborhood-based benchmarking and multilayer network topology.

**Results:** We observed many genes with altered expression in WF-exposed tissues were also among differentially expressed genes (DEGs) in RSD tissues; for CB, AS, PE and LC there were 34, 27, 50 and 26 genes respectively. DEG analysis, using disease association networks, pathways, ontological analysis and protein-protein interaction sub-network suggest significant links between WF exposure and the development of CB, AS, PE and LC.

**Conclusions:** Our network-based analysis and investigation of the genetic links of WFs and RSDs confirm a number of genes and gene products are plausible participants in RSD development. Our results are a significant resource to identify causal influences on the development of RSDs, particularly in the context of WF exposure.

## 1. Introduction

Welding is the process of joining metal at their contacting surface by using high temperature melting. It can very hazardous because it exposes the welder and others nearby to a variety of gases, WFs and radiant energy. Among the most dangerous WF components [1] are metallic oxides, fluorides and silicates including those of beryllium, aluminum, cadmium, chromium, copper, iron, lead, manganese, magnesium, nickel, vanadium, and zinc [2]. A known risk associated with large scale and long term WF exposure is the development of RSDs [1, 3]. The human respiratory system consists of many delicate tissues that transport air into the lungs for rapid uptake of oxygen in blood while eliminating carbon dioxide. The same exposed cell-rich surfaces that facilitate oxygen transport and uptake to the bloodstream also facilitate uptake of noxious gasses. In addition, such toxic chemicals can directly damage cells lining the respiratory system. This predisposes the respiratory system to develop RSDs such as asthma (AS), chronic bronchitis (CB), pulmonary edema (PE) and even lung cancer (LC) [4]. How WF and other pollutants do this is unclear, as indeed are the influences of many pathogens on lung tissues. One way to investigate the WF influences in causing or promoting such ill-health is to investigate gene expression effects of WFs on respiratory system cells and tissues and compare this with gene expression changes seen in tissues affected by CB, AS, PE and LC.

CB reduces the functionality of lung through damage to the cilia of the breathing passages and obstructs airflow inside the lung [5]. Iron, aluminum and magnesium oxide WFs may have particularly strong effects on this disease [6]. Asthma is a pulmonary system disorder that narrows and inflames the airways and is characterized by episodic wheezing, shortness of breath, chest tightness and coughing. It is often triggered by allergen exposure (which trigger auto-immune responses) and air pollutants [7]. Nitrogen oxides, carbon monooxide and phosgene in WFs are components particularly linked to asthma [6]. PE is a chronic lung condition characterized by fluid buildup in the lung, leading to serious breathing problems, coughing and chest pain [8]. Many irritants can drive PE but nickel, beryllium and cobalt oxides of the WFs have been linked to PE as well as LCs [6]. The latter, including both small cell and non-small cell lung carcinomas, are some of the deadliest types of cancer, and together are a major reported cause of death worldwide [9]. Most cases of LC are linked to tobacco smoke, though genetic factors and air pollutants play important roles. Such pollutants include many of the components of WFs, which can be taken up into, and affect, pulmonary tissues [6].

In this study, we developed a systematic and quantitative network-based approach to investigate how gene expression changes seen in WF exposure may interact with RSDs. Thus, we studied the gene expression in RSD affected tissues, namely CB, AS, PE and LC. To find the effects of WFs on gene expression in RSDs, we analyzed differentially expressed genes (DEGs) associated with these diseases (found by comparing gene expression in WF and RSD affected tissues with gene expression in controls), disease association network, signaling and ontology pathways, and protein-protein interaction networks. We also examined the validity of our study by employing the gold benchmark databases dbGAP and OMIM.

## 2. Materials and methods

### 2.1. Datasets employed in this study

To understand the genetic effects of WFs on RSDs at the molecular level, we analyzed gene expression microarray data. In this study, we employed gene expression microarray data from the National Center for Biotechnology Information (NCBI) (http://www.ncbi.nlm.nih.gov/geo/). We analyzed 5 different datasets with accession numbers GSE62384, GSE22148, GSE69683, GSE68610 and GSE10072 [10, 11, 12, 13, 14]. The WFs dataset (GSE62384) is a microarray data of fresh welding fumes influence on upper airway epithelial cells (RPMI 2650). This data is collected from the people with spark-generated welding fumes at high (760 g/m3) and low (85 g/m3) concentrations. The donors inhaled welding fumes for 6 hours continuously, followed by zero hours or four hours post-exposure incubation. The CB dataset (GSE22148) is taken from the 480 ex-smokers and stage 2 - 4 chronic obstructive pulmonary disease patients from the United Kingdom. The asthma dataset (GSE69683) is from microarray data collected from severe and moderate asthmatic patients and healthy subjects. The PE dataset (GSE68610) is an Affymetrix gene expression array of 25 acute lung injury patients. The LC dataset (GSE10072) is a microarray data where RNA was collected from 28 current smokers, 26 former smokers and 20 never smokers. In this data, final gene expression values have been reported by comparing between 58 tumor and 49 non-tumor lung tissues. The summarized description of the datasets are shown in table 1.

**Table 1:** Datasets description with GEO accession, GEO platform, data cells and number of simples.

| Sl. | Disease name | GEO accession | GEO platform | Data cells | Number of samples |  |
| --- | --- | --- | --- | --- | --- | --- |
|  |  |  |  |  | Case | Healthy |
| 1. | Welding fumes (WFs) | GSE62384 | GPL10558 | Nasal epithelial | 18 | 06 |
| 2. | Chronic bronchitis (CB) | GSE22148 | GPL570 | Sputum | 73 | 70 |
| 3. | Asthma (AS) | GSE69683 | GPL13158 | Blood | 87 | 411 |
| 4. | Pulmonary edema (PE) | GSE68610 | GPL571 | Alveolar epithelial | 17 | 08 |
| 5. | Lung cancer (LC) | GSE10072 | GPL96 | Lung | 58 | 49 |

### 2.2. Overview of the analytical approach

We applied a systematic and quantitative framework to investigate the genetic effects of WFs to the development of the RSDs by using several sources of available microarray datasets. The graphical representation of this approach is shown in figure 1. This approach uses gene expression microarray data to identify dysregulated genes, and identifies commonly dysregulated genes for each RSD with WFs. Further, these common dysregulated genes are employed to analyze signaling pathway, gene ontology (GO) and protein-protein interaction (PPI) sub-networks. Gold benchmark data is also used in this approach to verify the validation of our study.

**Figure 1:**
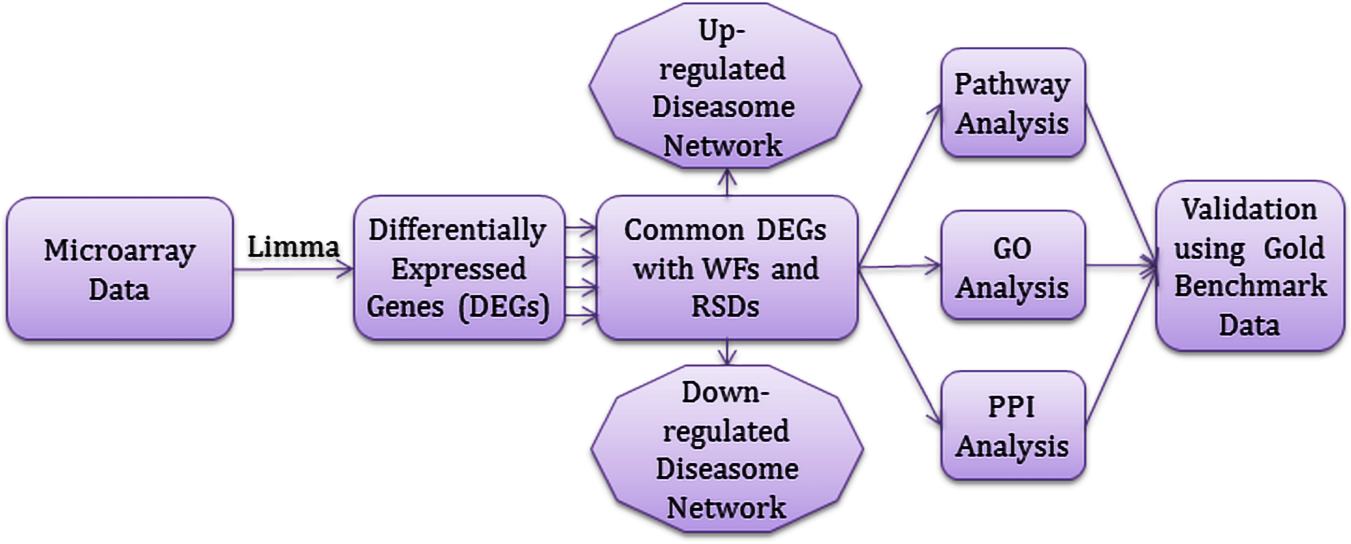
Flow-diagram of the analytical approach used in this study.

### 2.3. Analysis methods

Microarray based gene expression analysis is a global and sensitive method to find and quantify the human disorders at the molecular level [15]. We used these technologies to analyze the gene expression profiles of CB, AS, PE and LC to find the genetic effects of WFs on the development of respiratory system diseases. To make uniform mRNA expression data of different platforms and to avoid the problems of experimental systems, we normalized the gene expression data by using the Z-score transformation (*Z*_*ij*_) for each RSD gene expression profile using 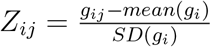,

where *SD* denotes the standard deviation, *g*_*ij*_ denotes the value of the gene expression *i* in sample *j*. After this transformation gene expression values of different diseases at different platforms can be directly compared. We applied two conditions for t-test statistic. We applied unpaired T-test to find differentially expressed genes of each disease over control data and selected significant dysregulated genes. We have chosen a threshold of at least 1 *log*_2_ fold change and a *p*-value for the t-tests of *<*= 1 * 10^−2^.

We employed neighborhood-based benchmark and the multilayer topological methods to find gene-disease associations. We constructed gene-disease network (GDN) using the gene-disease associations, where the nods in the network represent either gene or disease. This network can also be recognized as a bipartite graph. The primary condition for a disease to be connected with other diseases in GDN is they should share at least one or more significant dysregulated genes. Let *D* is a specific set of diseases and *G* is a set of dysregulated genes, gene-disease associations attempt to find whether gene *g ∈ G* is associated with disease *d ∈ D*. If *G*_*i*_ and *G*_*j*_, the sets of significant dysregulated genes associated with diseases *D*_*i*_ and *D*_*j*_ respectively, then the number of shared dysregulated genes 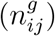 associated with both diseases *D*_*i*_ and *D*_*j*_ is as follows [15]:

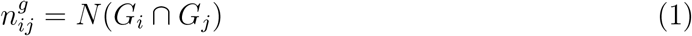

The common neighbours are the based on the Jaccard Coefficient method, where the edge prediction score for the node pair is as [15, 16]:

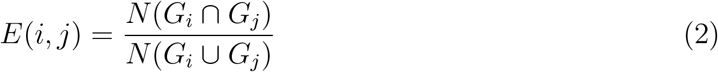

where *G* is the set of nodes and *E* is the set of all edges. We used R software packages ”comoR” [17] and ”POGO” [18] to cross check the genes-diseases associations.

To investigate the molecular determinants of WFs on several RSDs, we analyzed pathway and gene ontology using Enrichr (https://amp.pharm.mssm.edu/Enrichr/). We used KEGG, WikiPathways, Reactome and BioCarta databases for analyzing signaling pathway [19, 20, 21, 22]. We used GO Biological Process and Human Phenotype Ontology databases for ontological analysis [23, 24]. We also constructed a protein-protein interaction sub-network for each RSD with medium confidence of score 0.4, using the STRING database (https://string-db.org), a biological database and web resource of known and predicted protein-protein interactions. Furthermore, we examined the validity of our study by employing two gold benchmark databases OMIM (https://www.omim.org) and dbGAP (https://www.ncbi.nlm.nih.gov/gap).

## 3. Results

### 3.1. Gene expression analysis

We analyzed the gene expression microarray data from the National Center for Biotechnology Information (NCBI) (http://www.ncbi.nlm.nih.gov/geo/) for investigating the genetic effect of WFs to the development of RSDs. We observed that WF exposure results in 903 DEGs for which the adjusted *p <*= .01 and *|logFC| >*= 1. The DEGs of WFs contain 392 genes showing up and 511 with down-regulated expression. Similarly, we identified the most significant dysregulated genes for each RSD after applying statistical analysis. In CB We identified 678 DEGs (463 up and 215 down), in asthma 602 DEGs (297 up and 305 down), in PE 759 DEGs (404 up and 355 down) and in LC 890 DEGs (673 up and 217 down). We also performed cross-comparative analysis to find the DEGs common between the WF dataset and each RSD dataset. We observed that WFs shares a large number of DEGs with the other datasets, specifically 34, 27, 50 and 26 for CB, asthma, PE and LC respectively. To investigate the significant associations for DEGs from the RSDs with the WF datasets, we built two separate gene-disease association networks for up and down-regulated genes, centered on the WFs as shown in figure 2 and 3. The essential condition for two diseases to be associated with each is they should have at least one or more common genes in between them [25, 26]. Notably, 2 significant genes, PMAIP1 and SEC24A are commonly differentially expressed among WFs, PE, AS and LC; 6 genes, FGFR3, ID3, PROS1, AK4, TOX3 and MTHED2 are common to WFs, PE and LC. Similarly, PECR, ALDH3A2 and TPD52L1 are commonly differentially expressed in WFs, PE and CB; one gene HSD17BB is dysregulated among WFs, asthma and CB.

**Figure 2:**
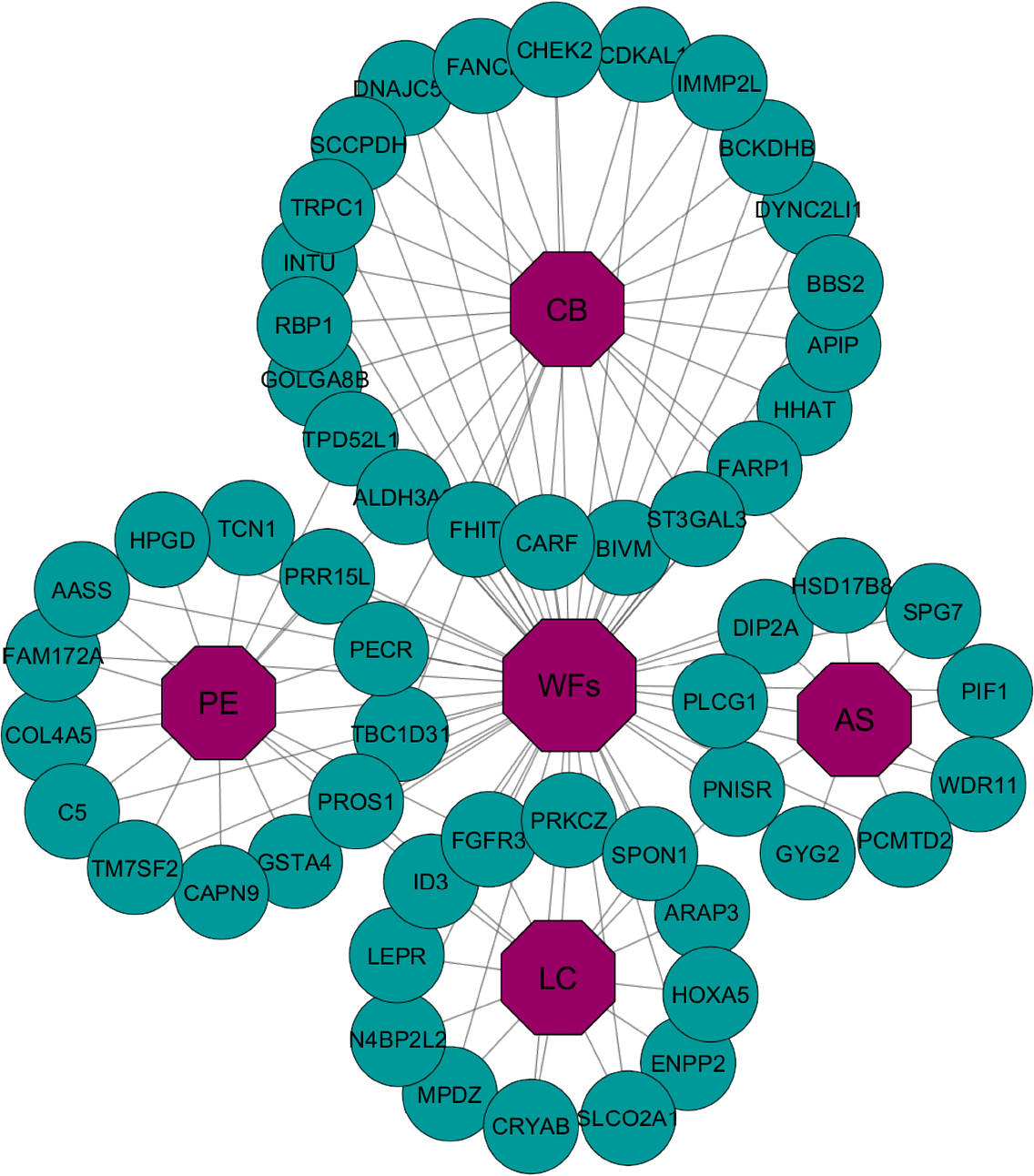
Gene-disease association network of welding fumes (WFs) with chronic bronchitis (CB), asthma (AS), pulmonary edema (PE) and lung cancer (LC). Octagon-shaped violet colored nodes represent different RSDs, and dark-cyan colored round-shaped nodes represent commonly up-regulated genes for WFs with the other RSDs. A link is placed between a disease and a gene if mutations in that gene lead to the specific disease.

**Figure 3:**
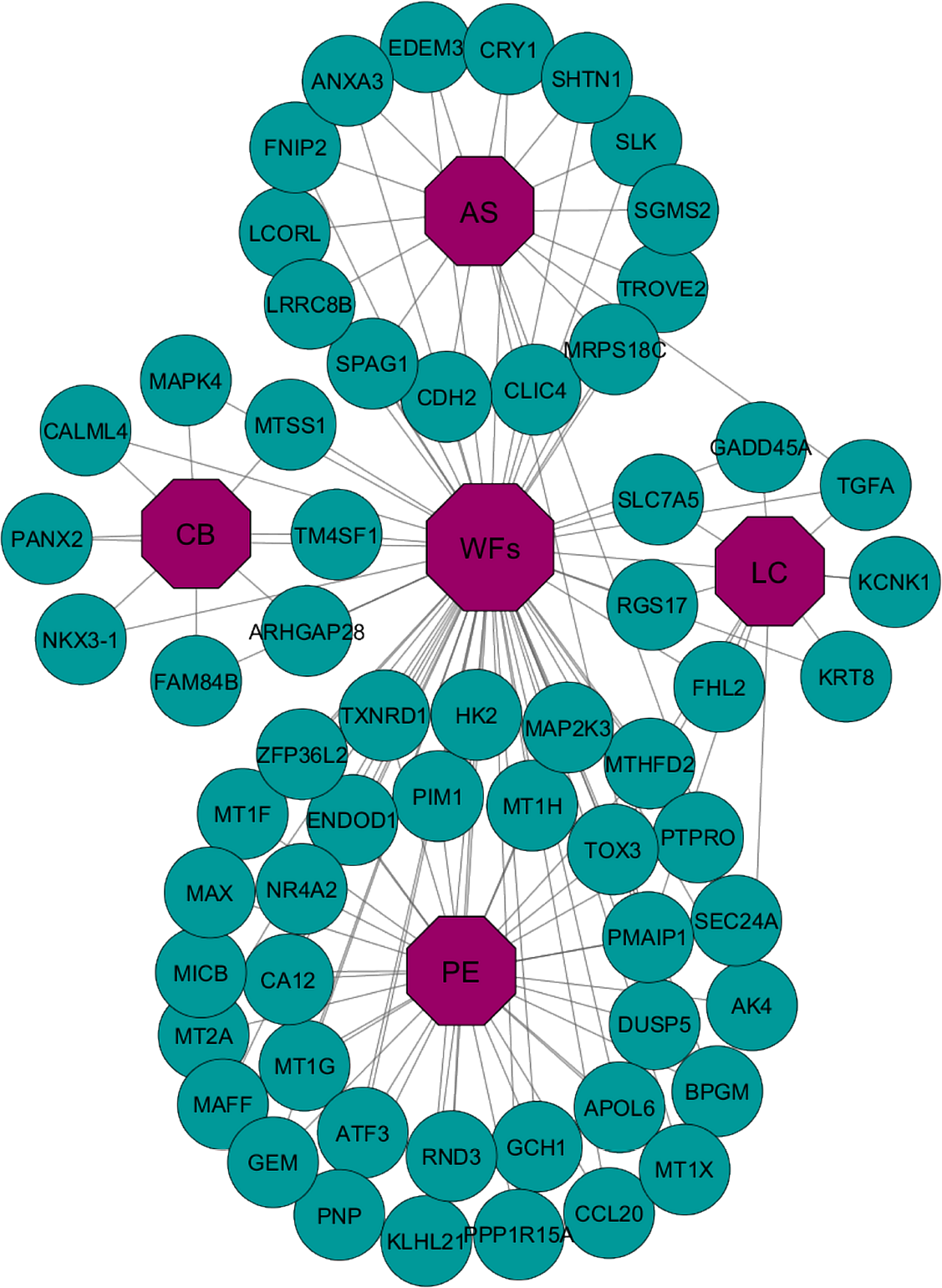
Gene-disease association network of welding fumes (WFs) with chronic bronchitis (CB), asthma (AS), pulmonary edema (PE) and lung cancer (LC). Octagon-shaped violet colored nodes represent different RSDs, and dark-cyan colored round-shaped nodes represent commonly down-regulated genes for WFs with the other RSDs. A link is placed between a disease and a gene if mutations in that gene lead to the specific disease.

### 3.2. Pathway and functional association analysis

Pathways are the important key to understand the reactions of an organism for its internal changes. The pathway-based analysis is a modern technique to understand how different complex diseases are related to each other by underlying molecular or biological mechanisms [27, 28]. We analyzed pathways of the commonly dysregulated genes of WFs and each RSD using Enrichr, a comprehensive web-based tool for analyzing gene set enrichment [29]. Signaling pathways of the commonly differentially expressed genes in between WFs and each RSD were analyzed using four global databases includes KEGG, WikiPathways, Reactome and BioCarta. We collected and combined pathways from the mentioned four databases and selected the most significant pathways of each RSD after several steps of statistical analysis.

Four significant pathways associated with CB are shown in table 2. Among these pathways, ’Integrated Pancreatic Cancer Pathway’ contains some proteins that are known as important in human breast cancer development [30, 31]. The pathway ’Assembly of the motile cilium’ is only known in cilia functions such as respiratory cilia [32]. ’Methionine salvage’ is a pathway that performs six reactions in the human body and is responsible for the recycling of sulfur in the respiratory system [33]. The pathway ’Glyoxylate metabolism and glycine degradation’ is responsible for overproduction of oxalate in human liver and lung [34].

**Table 2:** Signalling pathways associated with significantly commonly altered genes of the CB with WFs.

| Pathway | Genes in the pathway | Adjusted p-value |
| --- | --- | --- |
| Integrated pancreatic cancer pathway | CHEK2, MAPK4 | 3.98E-02 |
| Assembly of the motile cilium | BBS2, DYNC2LI1 | 3.80E-02 |
| Methionine salvage pathway | APIP | 9.86E-03 |
| Glyoxylate metabolism and glycine degradation | BCKDHB | 4.05E-02 |

The top five significant pathways associated with AS are shown in table 3. Among these pathways, ’Non-small cell lung cancer’ is linked to development of adeno-squamous cell and large-cell carcinoma types that are responsible for approximately 75% of all lung cancer [35, 36]. ’IL-1 Signaling Pathway’ is responsible to control pro-inflammatory responses, including inflammasome activity tissues [37]. ’Glycogen synthesis’ is responsible for several reactions to produce glycogen in liver, muscle, lung and other tissues; glycogen serves as a major stored short-term fuel in muscle and other high metabolism cells [38]. ’Constitutive Signaling by EGFRvIII’ is linked to the general development of tumors. ’Signaling by EGFRvIII in Cancer’ is more responsible for cancer in several cell types found in the respiratory system.

**Table 3:** Signalling pathways associated with significantly commonly altered genes of the AS with WFs.

| Pathway | Genes in the pathway | Adjusted p-value |
| --- | --- | --- |
| Non-small cell lung cancer | TGFA, PLCG1 | 2.40E-03 |
| IL-1 signaling pathway | PLCG1 | 4.58E-02 |
| Glycogen synthesis | GYG2 | 1.93E-02 |
| Constitutive signaling by EGFRvIII | PLCG1 | 1.93E-02 |
| Signaling by EGFRvIII in cancer | PLCG1 | 1.93E-02 |

Five significant pathways associated with PE are indicated in table 4. These include ’Central carbon metabolism in cancer’ which includes three transcription factors HIF-1, c-MYC and p53 that influence the regulate tumor cell growth [39]. ’Small cell lung cancer’ pathway relates to a highly aggressive neoplasm responsible for approximately 25% of all lung cancer [40]. ’TNF signaling pathway’ is involved in the control of inflammation, immunity and cell survival. ’One carbon pool by folate’ is pathway linked to progression of several forms of cancer [41]. ’Response to metal ions’ pathway may mediate cell toxicity effects of heavy metals such as zinc, copper, and iron [42].

**Table 4:** Signalling pathways associated with significantly commonly altered genes of the PE with WFs.

| Pathway | Genes in the pathway | Adjusted p-value |
| --- | --- | --- |
| Central carbon metabolism in cancer | FGFR3, HK2 | 1.17E-02 |
| Small cell lung cancer | MAX, COL4A5 | 1.89E-02 |
| TNF signaling pathway | MAP2K3, CCL20 | 2.98E-02 |
| One carbon pool by folate | MTHFD2 | 4.79E-02 |
| Response to metal ions | MT2A, MT1F, MT1G | 3.27E-02 |

The top five significant pathways associated with LC are shown in table 5. Among these pathways, ’Central carbon metabolism in cancer’ is responsible for three transcription factors HIF-1, c-MYC and p53 that can coordinate regulation of tumor and cancer in cell [39]. ’Bladder Cancer’ is responsible for developing several carcinomas in urinary tract, renal pelvis, ureter and bladder [43]. ’Signaling by activated point mutants of FGFR3’ is responsible for several cancer including breast, prostate, bladder, cervical, neck and head [44]. ’Signaling by FGFR3 fusions in cancer’ is specifically responsible for lung and bladder cancer [45]. ’TGF-beta receptor signaling in EMT’ is responsible for the development of tumors in the early stage that can bring cancer in cells [46].

**Table 5:** Signalling pathways associated with significantly commonly altered genes of the LC with WFs.

| Pathway | Genes in the pathway | Adjusted p-value |
| --- | --- | --- |
| Central carbon metabolism in cancer | SLC7A5, FGFR3 | 3.16E-03 |
| Bladder cancer | FGFR3 | 3.81E-02 |
| Signaling by activated point mutants of FGFR3 | FGFR3 | 1.49E-02 |
| Signaling by FGFR3 fusions in cancer | FGFR3 | 1.24E-02 |
| TGF-beta receptor signaling in EMT | PRKCZ | 1.98E-02 |

### 3.3. Gene ontological analysis

The Gene Ontology (GO) refers to a universal conceptual model for representing gene functions and their relationship in the domain of gene regulation. It is constantly expanded by accumulating the biological knowledge to cover regulation of gene functions and the relationship of these functions in terms of ontology classes and semantic relations between classes [47]. GO of the commonly dysregulated genes for each RSD and WFs were analyzed using two databases including GO Biological Process and Human Phenotype Ontology. We collected and concatenated gene ontologies from mentioned two databases, and performed several statistical analysis to identify the most significant ontologies of each respiratory system diseases. Notably, we found 12, 13, 10 and 15 gene ontology terms are associated with the CB, AS, PE and LC respectively as shown in table 6–9.

**Table 6:** Ontological pathways associated with significantly commonly altered genes of the CB with WFs.

| GO Term | Pathway | Genes in the pathway | Adjusted p-value |
| --- | --- | --- | --- |
| GO:2001233 | Regulation of apoptotic signaling pathway | TPD52L1, NKX3-1 | 8.37E-04 |
| GO:0060562 | Epithelial tube morphogenesis | MTSS1, NKX3-1 | 1.52E-03 |
| GO:0050678 | Regulation of epithelial cell proliferation | MTSS1, NKX3-1 | 6.45E-03 |
| GO:0046467 | Membrane lipid biosynthetic process | ALDH3A2, SCCPDH | 4.55E-03 |
| GO:0051225 | Spindle assembly | GOLGA8B, CHEK2 | 7.89E-03 |
| GO:2001235 | Positive regulation of apoptotic signaling pathway | TPD52L1, NKX3-1 | 7.15E-03 |
| HP:0100631 | Neoplasm of the adrenal gland | CHEK2 | 2.61E-02 |
| HP:0002858 | Meningioma | CHEK2 | 2.77E-02 |
| HP:0007707 | Congenital primary aphakia | BBS2 | 2.93E-02 |
| HP:0100641 | Neoplasm of the adrenal cortex | CHEK2 | 1.64E-02 |
| HP:0002476 | Primitive reflexes | ST3GAL3 | 1.48E-02 |
| HP:0012126 | Stomach cancer | CHEK2 | 1.48E-02 |

**Table 7:** Ontological pathways associated with significantly commonly altered genes of the AS with WFs.

| GO Term | Pathway | Genes in the pathway | Adjusted p-value |
| --- | --- | --- | --- |
| GO:2000323 | Negative regulation of glucocorticoid receptor signaling pathway | CRY1 | 9.07E-03 |
| GO:0031334 | Positive regulation of protein complex assembly | PMAIP1, FNIP2 | 5.94E-03 |
| GO:1904018 | Positive regulation of vasculature development | ANXA3, PLCG1 | 8.17E-03 |
| GO:0010634 | Positive regulation of epithelial cell migration | ANXA3, PLCG1 | 4.25E-03 |
| GO:0032147 | Activation of protein kinase activity | SLK, TGFA, PLCG1 | 3.37E-03 |
| GO:0043067 | Regulation of programmed cell death | SLK, PMAIP1, DIP2A | 4.98E-03 |
| HP:0002492 | Abnormality of the corticospinal tract | SPG7 | 1.29E-02 |
| HP:0004469 | Chronic bronchitis | SPAG1 | 1.55E-02 |
| HP:0012261 | Abnormal respiratory motile cilium physiology | SPAG1 | 2.44E-02 |
| HP:0200073 | Respiratory insufficiency due to defective ciliary clearance | SPAG1 | 1.16E-02 |
| HP:0012384 | Rhinitis | SPAG1 | 2.44E-02 |
| HP:0000458 | Anosmia | WDR11 | 2.82E-02 |
| HP:0012387 | Bronchitis | SPAG1 | 3.58E-02 |

**Table 8:**
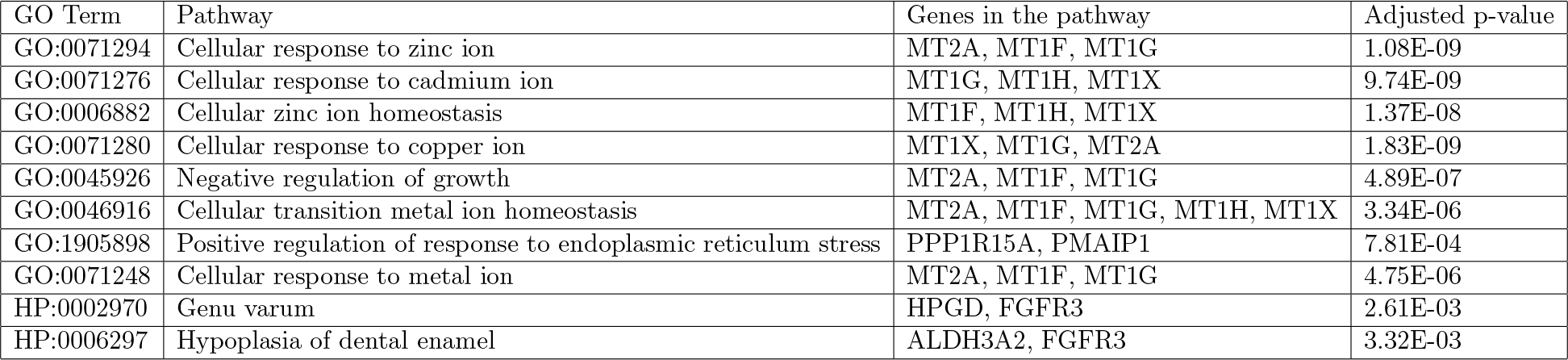
Ontological pathways associated with significantly commonly altered genes of the PE with WFs.

| GO Term | Pathway | Genes in the pathway | Adjusted p-value |
| --- | --- | --- | --- |
| GO:0071294 | Cellular response to zinc ion | MT2A, MT1F, MT1G | 1.08E-09 |
| GO:0071276 | Cellular response to cadmium ion | MT1G, MT1H, MT1X | 9.74E-09 |
| GO:0006882 | Cellular zinc ion homeostasis | MT1F, MT1H, MT1X | 1.37E-08 |
| GO:0071280 | Cellular response to copper ion | MT1X, MT1G, MT2A | 1.83E-09 |
| GO:0045926 | Negative regulation of growth | MT2A, MT1F, MT1G | 4.89E-07 |
| GO:0046916 | Cellular transition metal ion homeostasis | MT2A, MT1F, MT1G, MT1H, MT1X | 3.34E-06 |
| GO:1905898 | Positive regulation of response to endoplasmic reticulum stress | PPP1R15A, PMAIP1 | 7.81E-04 |
| GO:0071248 | Cellular response to metal ion | MT2A, MT1F, MT1G | 4.75E-06 |
| HP:0002970 | Genu varum | HPGD, FGFR3 | 2.61E-03 |
| HP:0006297 | Hypoplasia of dental enamel | ALDH3A2, FGFR3 | 3.32E-03 |

**Table 9:** Ontological pathways associated with significantly commonly altered genes of the LC with WFs.

| GO Term | Pathway | Genes in the pathway | Adjusted p-value |
| --- | --- | --- | --- |
| GO:1903708 | Positive regulation of hemopoiesis | N4BP2L2, HOXA5 | 4.18E-05 |
| GO:0009132 | Nucleoside diphosphate metabolic process | AK4 | 9.96E-03 |
| GO:0045647 | Negative regulation of erythrocyte differentiation | HOXA5 | 8.72E-03 |
| GO:2000665 | Regulation of interleukin-13 secretion | PRKCZ | 8.72E-03 |
| GO:0044320 | Cellular response to leptin stimulus | LEPR | 9.96E-03 |
| GO:0045837 | Negative regulation of membrane potential | PMAIP1 | 8.72E-03 |
| GO:2000394 | Positive regulation of lamellipodium morphogenesis | ENPP2 | 9.96E-03 |
| GO:0050730 | Regulation of peptidyl-tyrosine phosphorylation | ENPP2, PRKCZ | 5.14E-03 |
| GO:0032754 | Positive regulation of interleukin-5 production | PRKCZ | 9.96E-03 |
| GO:0043065 | Positive regulation of apoptotic process | GADD45A, PMAIP1 | 6.47E-03 |
| HP:0000975 | Hyperhidrosis | SLCO2A1, FGFR3 | 5.38E-03 |
| HP:0000522 | Alacrima | FGFR3 | 1.24E-02 |
| HP:0010662 | Abnormality of the diencephalon | LEPR | 1.86E-02 |
| HP:0000495 | Recurrent corneal erosions | FGFR3 | 1.49E-02 |
| HP:0001413 | Micronodular cirrhosis | KRT8 | 1.86E-02 |

### 3.4. Protein-protein interaction analysis

Protein-protein interaction network (PPIN) is the graphical representation of the physical connection of proteins in the cell. Protein-protein interactions (PPIs) are essential to every molecular and biological process in a cell, so PPIs is crucial to understand cell physiology in disease and healthy states [48]. We have used STRING database to analyze and construct protein-protein interaction subnetworks of the significantly commonly dysregulated genes of each RSD. We have clustered into four different groups of protein-protein interactions of four RSDs as shown in figure 4.

**Figure 4:**
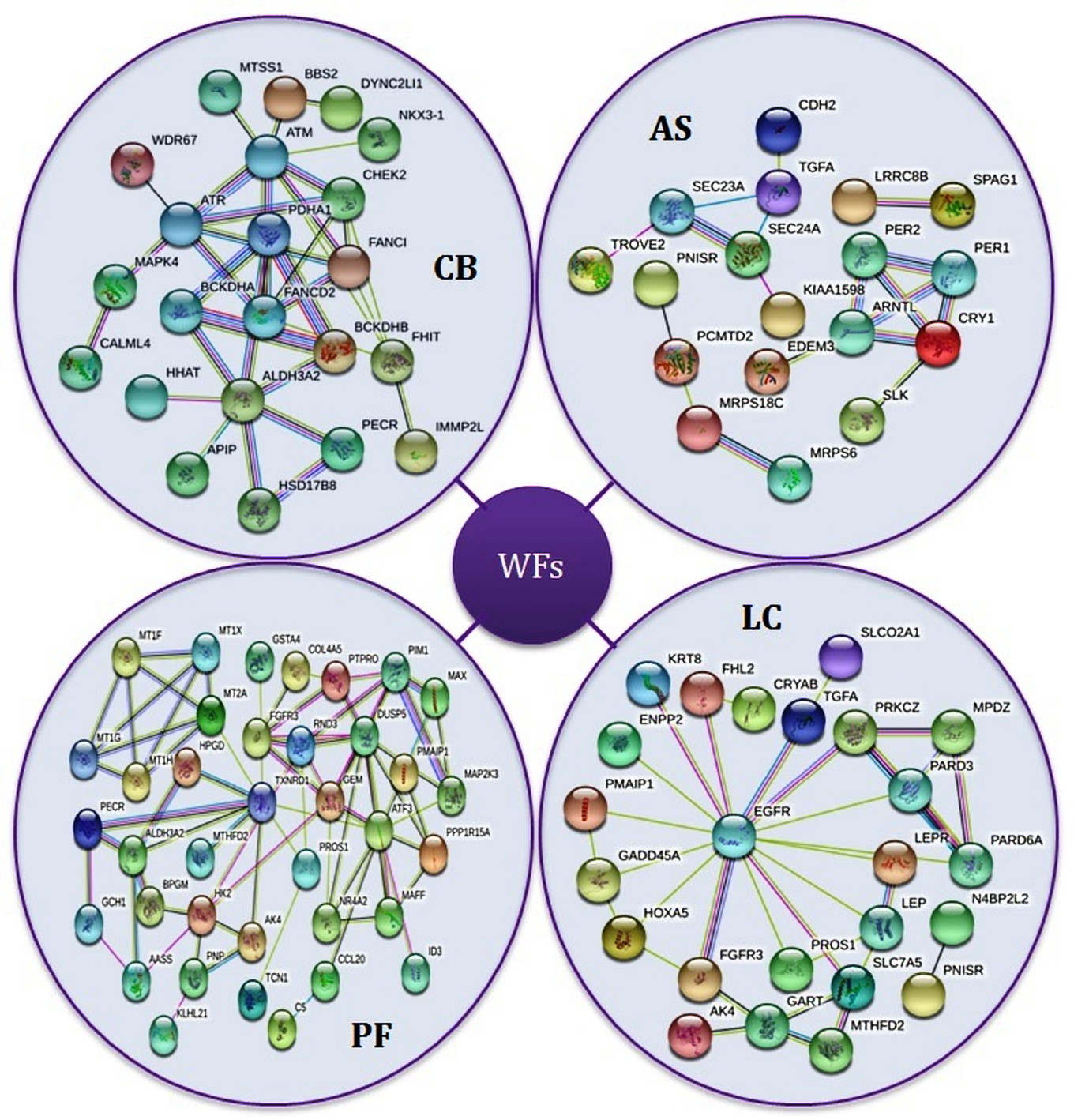
Protein-protein interaction network of RSDs using STRING.

## 4. Discussion

WFs are well understood to have toxic effects on lung tissues, so that chronic exposure to WFs increases the risk of developing AS, CB, PE and LC [49]. Nevertheless, how WFs may drive lung changes that predispose to these diseases is not well understood. Zeidler-Erdely have studied WF effects on lungs of mice [50] which found that mouse lung transcriptomes were altered by WF exposure induced expression of immunomodulatory chemokines. However, this was more acute exposure than chronic. WF influences on respiratory diseases were studied by Jnson et al who examined gene expression in sputum cells of welders with respiratory problems [51]. The gene clusters identified included inflammatory response, defence response and Wounding response and reported mostly secreted cytokines. Clear evidence of carcinogenic effects of WF in lung were found by Li et al. [52] observed that WF exposure altered methylation patterns of tumour suppressor genes were altered. However, there is little other direct evidence regarding how chronic effects of WF on lungs can encourage respiratory diseases and cancer. We can study this by examining gene expression effects of WF in lung tissues and see how gene pathway activation might be pathogenic; we have used an analogous approach in a previous study of WF exposure and neurological diseases [52].

With this in mind, we investigated WF exposure gene regulation data with that of RSDs to identify the common gene networks they have in common, including signaling pathways, gene expression ontologies and protein-protein interaction sub-networks. For the purpose of the present study, we analyzed gene expression microarray data from WFs, CB, AS, PE, LC and control datasets. We identified a large number of of significant DEGs in common between WFs and RSDs which suggests that WFs may induce gene expression mediated effects in lung cells that increase the incidence and/or severity of RSDs. We constructed two separate gene-disease association networks for up and down-regulated genes showed a strong evidence that WFs are linked to the development of RSDs as shown in Figure 2 and 3. The pathway-based analysis is useful to understand how different complex diseases are related to each other by underlying molecular or biological mechanisms. We identified significant pathways common to the dysregulated geneset of each RSD and the WF data. Similarly, gene expression ontologies and protein-protein interaction sub-networks of the commonly differentially expressed genes suggest that WFs may be a major stimulus for progression of several RSDs.

We have used the gold benchmark databases (dbGAP and OMIM) to verify the outcome of our study and observed that there are a number of shared genes between the WFs and RSDs as shown in figure 5. We collected disease and genes names from OMIM Disease, OMIM Expanded and dbGap databases using the observed altered genes seen in WF exposure for cross checking the validity of our study. We concatenated the list of diseases as well as genes from the mentioned three databases and selected only RSDs after several steps of statistical analysis. Interestingly, we found our selected four RSDs among the list of collected diseases as shown in figure 5. Moreover, we found our identified genes in figure 5 had been shown in other studies to be associated with disease progression in RSDs. Specifically, Lamontagne M. et al. have shown a link between RAB4B and CB [53]; D’Anna C. at el. found ANXA5 to be associated to CB incidence [54]; Holt RJ et al. showed POU2F1 to be linked to AS [55]; Sinn H. et al. found FOS to be linked to AS incidence [56]; Imboden M. et al. had shown SVEP1 to be associated with AS [57]; Boor P. et at. found C5 to be linked to PE [58]; Desai J. et al. have identified a link between KIF21A and PE [59]; Clark AM. et al. found an association between MAP3K8 and LC [60]; Mangia A. et al. found SLC9A3R1 to be associated to LC incidence [61]. These genes and their products are thus strong candidate mediators for the deleterious effects of WFs and lung that predispose welders to develop CB, AS, PE and LC.

**Figure 5:**
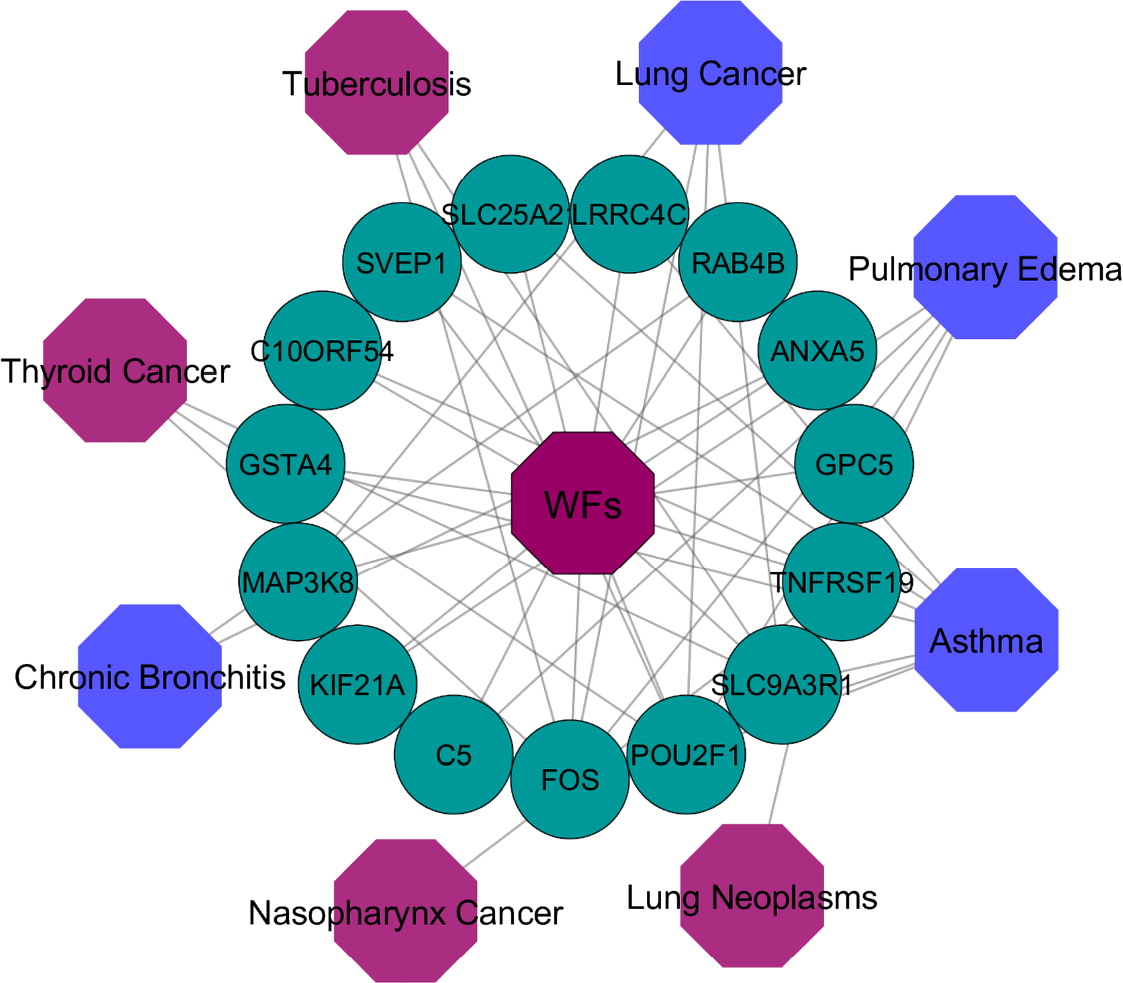
Gene-disease association network of welding fumes (WFs) with several RSDs. Violet colored octagon-shaped nodes represent different RSDs, blue colored octagon-shaped nodes represent our selected four RSDs and round-shaped dark-cyan colored nodes represent differentially expressed genes for WFs. A link is placed between a disease and a gene if mutations in that gene lead to the specific disease.

In table 6 and figure 4 we delineate a number of pathways we identified as important our WF-diseasome analyses. Regulation of apoptotic signaling pathway relates to regulation of programmed cell death that is essential for normal functions of tissues; this can be increased when tissue damage is occurring. Inhibiting such apoptosis may better preserve tissue integrity, but it may leave tissues vulnerable to cancer transformation. Cell proliferation is similarly crucial in tissue response to damage, particularly when high levels of tissue damage occurs, and we identified several pathways that influence this, including Regulation of epithelial cell proliferation, Membrane lipid biosynthetic process and Spindle assembly. In several cases there are genes that are associated with multiple pathways, such as NKX3-1, which belongs to homeobox genes [62] that regulate body morphology and renewal at a high level, i.e., by influencing expression of many genes. Indeed, this gene and MTSS1 (metastasis suppressor gene and Src-type tyrosine kinase inhibitor [63]) form part of the Epithelial tube morphogenesis pathway. Our study has identified a number of hub protein genes that are important high level controllers that will regulate complex tissue changes. NKX3-1 is also important in pluripotent stem cell maintenance [64]. It is notable that many of these genes and pathways are implicated in cancer-related functions, which might be expected as chronic damage caused by WFs leads to tissue repair functions that are also extensively utilised in cancer progression. The pathways identified are thus consistent with the tissue damage and healing seen in both the WFs and associated diseases, but also identify particular means in which WFs influence those diseases and in consequence may indicate high pathogenic potential for the genes involved.

## 5. Conclusions

In this study, we have considered gene expression microarray data from WFs, CB, AS, PE, and LC datasets to analyze and investigate the genetic effects of WFs on RSDs. We analyzed gene regulation, built gene-disease association networks, identified signaling pathways, identified gene expression ontologies and protein-protein interaction sub-networks of WFs and each RSDs. Our findings showed the nature of the association between WFs and RSDs at the molecular and cellular level. This types of study are likely to be useful for making gene-based evidence based recommendations about accurate disease prediction, identification and therapeutic treatments and may identify gene pathway elements that are of particular pathogenic important to thee lung diseases.

